# Philympics 2021: Prophage Predictions Perplex Programs

**DOI:** 10.1101/2021.06.03.446868

**Authors:** Michael J. Roach, Katelyn McNair, Sarah K. Giles, Laura Inglis, Evan Pargin, Simon Roux, Przemysław Decewicz, Robert A. Edwards

## Abstract

Most bacterial genomes contain integrated bacteriophages—prophages—in various states of decay. Many are active and able to excise from the genome and replicate, while others are cryptic prophages, remnants of their former selves. Over the last two decades, many computational tools have been developed to identify the prophage components of bacterial genomes, and it is a particularly active area for the application of machine learning approaches. However, progress is hindered and comparisons thwarted because there are no manually curated bacterial genomes that can be used to test new prophage prediction algorithms.

Here, we present a library of gold-standard bacterial genome annotations that include manually curated prophage annotations, and a computational framework to compare the predictions from different algorithms. We use this suite to compare all extant stand-alone prophage prediction algorithms to identify their strengths and weaknesses.

We provide a FAIR dataset for prophage identification, and demonstrate the accuracy, precision, recall, and f_1_ score from the analysis of seven different algorithms for the prediction of prophages. We discuss caveats and concerns in this analysis and how those concerns may be mitigated.

## Introduction

Bacteriophages (phages), viruses that infect bacteria, can be either temperate or virulent. Temperate phages may integrate into their bacterial host genome and the host-integrated phage genome is referred to as a prophage. Prophages are ubiquitous and may constitute as much as 20 percent of bacterial genomes (Casjens, 2003). Prophages replicate as part of the host bacterial genomes until external conditions trigger a transition into the virulent lytic cycle, resulting in replication and packaging of phages and typically the death of the host bacteria. Prophages generally contain a set of core genes with a conserved gene order that facilitate integration into the host genome, assembly of phage structural components, replication, and lysis of the host cell (Kang et al., 2017, Canchaya et al., 2003). As well as these core genes, phages can contain an array of accessory metabolic genes that can effect significant phenotypic changes in the host bacteria (Breitbart, 2012). For instance, many prophages encode virulence factors such as toxins, or they can encode fitness factors such as nutrient uptake systems (Brüssow et al., 2004). Lastly, most prophages encode a variety of super-infection exclusion mechanism to prevent concurrent phage infections, including restriction/modification systems, toxin/antitoxin genes, repressors, etc. (Calendar, 1988). The function of most prophage accessory genes remains unknown.

Core (pro)phage genes have long been used for identifying prophage regions. However, there are other unique characteristics that can distinguish prophages from their host genomes: bacterial genomes have a GC skew that correlates with direction of replication, and the insertion of prophages will generally disrupt this GC bias (Grigoriev, 1998). Transcript direction (Campbell, 2002) and length of prophage proteins have also proven to be useful metrics in predicting prophages (Akhter et al., 2012, Song et al., 2019), where phage genes are generally smaller and are oriented in the same direction (Dutilh et al., 2014). Likewise, gene density tends to be higher in phage genomes and intergenic space shorter (Amgarten et al., 2018, McNair et al., 2019).

Over the last two decades many prophage prediction tools have been developed, and they fall into two broad classes: (1) web-based tools where users upload a bacterial genome and retrieve annotations including PHASTER (Arndt et al., 2016), Prophage Hunter (Song et al., 2019), Prophinder (Lima-Mendez et al., 2008), PhageWeb (Sousa et al., 2018), and RAST (Aziz et al., 2008); and (2) command-line tools where users download a program and database to run the predictions locally (although some of these also provide a web interface for remote execution). In this work we focus on this latter set of tools (Table 1) because web-based tools typically do not handle the large numbers of simultaneous requests required to run comparisons across many genomes.

**Table 1:**
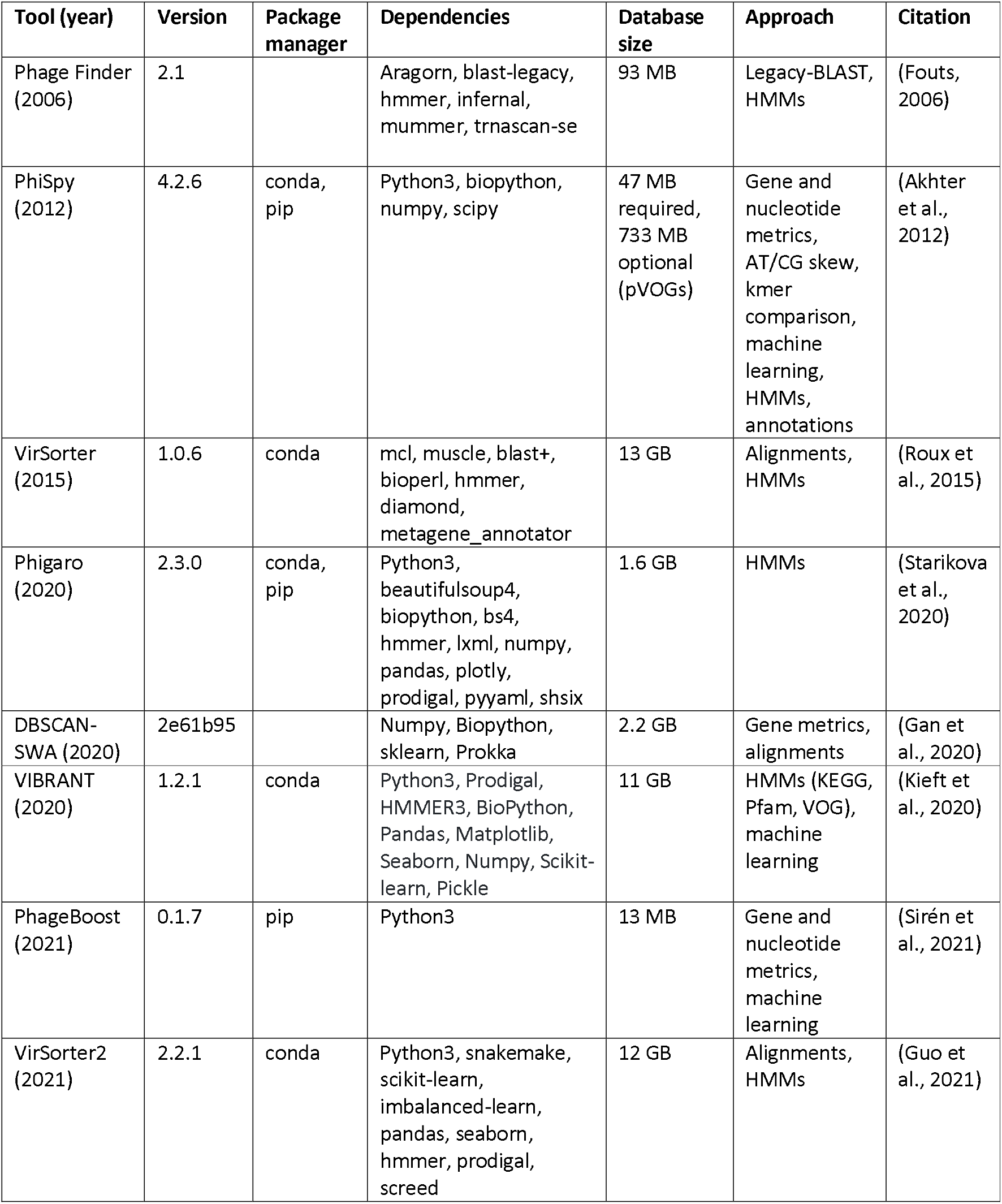
Prophage identification tools currently included in benchmarking framework

Despite the abundance of prophage prediction algorithms, there has never been either a set of reference genomes against which all tools can be compared, nor a unified framework for comparing those tools to identify their relative strengths and weaknesses or to identify opportunities for improvement. We generated a set of manually annotated bacterial genomes released under the FAIR principles (Findable, Accessible, Interoperable, and Reusable), and developed an openly available and accessible framework to compare prophage prediction tools.

## Methods

### Running the tools

To assess the accuracy of the different prophage prediction tools, a set of 49 gold-standard publicly available bacterial genomes with manually curated prophage annotations was generated. The genomes and prophage annotations currently included are available in Tables S1 and S2. The genomes are in GenBank format and file conversion scripts are included in the framework to convert those files to formats used by the different software. The tools that are currently included in the framework are outlined in Table 1. Snakemake (Köster and Rahmann, 2012) pipelines utilising conda (Anaconda Software Distribution. *Conda*. v4.10.1, April 2021) package manager environments were created for each tool to handle the installation of the tool and its dependencies, running of the analyses, output file conversion to a standardized format, and benchmarking of the run stage. Where possible, annotations from the GenBank files were used in the analysis to promote consistency between comparisons. Additional pipelines were created for running PhiSpy using the included training sets for the appropriate genera, and for running PhiSpy with pVOG (Grazziotin et al., 2017) HMMs and these are also available in the repository. DBSCAN-SWA was not able to consistently finish when using GenBank files as input, and instead the genome files in fasta format were used. Another pipeline was created to pool the results from each tool and some comparisons are illustrated in the included Jupyter notebook. Testing and development of the pipelines were conducted on Flinders University’s DeepThought HPC infrastructure. The final benchmarking analysis was performed on a stand-alone node consisting of dual Intel^®^ Xeon^®^ Gold 6242R processors (40 cores, 80 threads), 768 GB of RAM, and 58 TB of disk space. Each tool was executed on all genomes in parallel (one thread per job), with no other jobs running.

### Benchmark metrics

There are many potential ways to compare prophage predictions: For instance, is it more important

##### Box 1. Benchmark Metrics Used in this Analysis

Accuracy was calculated as the ratio of correctly labelled genes to all CDS features from the GenBank file.

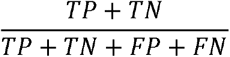

Precision was calculated as the ratio of correctly labelled phage CDS features to all predicted prophage CDS features

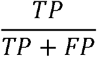

Recall was calculated as the ratio of correctly labelled prophage CDS features to all known prophage CDS features

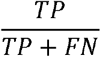

The f1 Score was calculated as the harmonic mean of Precision and Recall

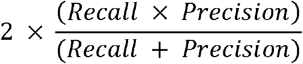

*Accuracy* provides an overall impression of correctness but is distorted by the vast difference in the numbers of prophage and non-prophage CDS features present in the genomes. The current gold-standard set includes 7,729 prophage proteins and 177,649 non-prophage proteins. Therefore, predicting everything as not coming from a prophage will result in an accuracy of 0.96. Similarly, identifying everything as coming from a prophage will result in high *Recall*, since that favours minimising false negatives. In contrast, *Precision* favours minimising false-positives and so only predicting very confident regions will result in high precision. The f1 Score is the most suitable for comparing predictions as it gives equal weighting to both precision and recall, and thus balances the unevenness inherent in this data.

to capture all prophage regions or minimise false positives? Is it more important to identify all the phage-encoded genes, or the exact locations of the attachment site core duplications (*attL* and *attR*)? The runtime and CPU time in seconds, peak memory usage and file write operations were captured by Snakemake for the steps running the prophage tools only (not for any file conversion steps before or after running each tool). The predictions were then compared to the gold standard annotations and the number of true positive (TP), true negative (TN), false positive (FP) and false negative (FN) gene labels were used to calculate the performance metrics. Each application marks prophages slightly differently, and therefore we used the designation of coding sequence (CDS) features as phage or not to assess prophage predictions.

### Adding new genomes

We developed the framework to simplify the addition of new genomes to the benchmarks. Each genome is provided in the standard GenBank format, and the prophages are marked by the inclusion of a non-standard flag for each genomic feature that indicates that it is part of a prophage. We use the qualifier */is_phage=”1”* to indicate prophage regions.

## Results and Discussion

### Software Compared

We compared the availability, installation, and results from ten different prophage prediction algorithms (Table 1). Two—ProphET (Reis-Cunha et al., 2019) and LysoPhD (Niu et al., 2019) —could not be successfully installed and were not included in the current framework (see below). The remaining eight PhiSpy (Akhter et al., 2012), Phage Finder (Fouts, 2006), VIBRANT (Kieft et al., 2020), VirSorter (Roux et al., 2015), Virsorter2 (Guo et al., 2021), Phigaro (Starikova et al., 2020), PhageBoost (Sirén et al., 2021), and DBSCAN-SWA (Gan et al., 2020) were each used to predict the prophages in 49 different manually curated microbial genomes.

Most of these programs utilize protein sequence similarity and HMM searches of core prophage genes to identify prophage regions. PhageBoost leverages a large range of protein features (such as dipeptide and tripeptide combinations) with a trained prediction model. PhiSpy was originally designed to identify prophage regions based upon seven distinct characteristics: protein length, transcript directionality, AT and GC skew, unique phage words, phage insertion points, optionally phage protein similarity and sequence similarity. DBSCAN-SWA likewise uses a range of gene metrics and trained prediction models to identify prophages.

Regardless of whether annotations are available, Virsorter2, Phigaro, and PhageBoost all perform *de novo* gene prediction with Prodigal (Hyatt et al., 2010) and VirSorter uses MetaGeneAnnotator (Noguchi et al., 2008) for the same purpose. VIBRANT can take proteins if they have ‘Prodigal format definition lines’ but otherwise performs predictions with Prodigal. PhageBoost can take existing annotations but this requires additional coding by the user. DBSCAN-SWA can take annotations or can perform gene predictions with Prokka (Seemann, 2014). PhiSpy takes an annotated genome in GenBank format and uses the annotations provided.

### Ease of installation

The prophage prediction packages Phigaro, PhiSpy, VIBRANT, VirSorter, and VirSorter2 are all able to be installed with conda from the Bioconda channel (Grüning et al., 2018), while Phispy, Phigaro, and PhageBoost can be installed with pip—the Python package installer. Phigaro, VIBRANT, VirSorter, and VirSorter2 require a manual one-time setup to download their respective databases. Phigaro uses hard-coded file paths for its database installation, either to the user’s home directory or to a system directory requiring root permissions. Neither option is ideal as it is impossible to have isolated versions or installations of the program, and it prevents updating the installation paths of its dependencies. For PhageBoost to be able to take existing annotations, a custom script was created to skip the gene prediction stage and run the program. Basic PhiSpy functionality is provided without requiring third-party databases. However, if the HMM search option is invoked, a database of phage-like proteins— e.g. pVOG (Grazziotin et al., 2017), VOGdb (https://vogdb.org), or PHROGS (Terzian P et al., 2021)—must be manually downloaded before it can be included in PhiSpy predictions. DBSCAN-SWA is not currently available on any package manager and must be pulled from GitHub, however all its dependencies are available via conda and it could easily be added in the future. All the above “manual” installation and setup steps are uncomplicated and are automatically executed by the Snakemake pipelines provided in the framework.

Phage Finder was last updated in 2006 and is not available on any package manager that we are aware of. The installation process is dated with the package scripts liberally utilising hard-coded file paths. The Snakemake pipeline for this package resolves this with soft links between the framework’s directory to the user’s home directory (where the package expects to be installed). The dependencies are available via conda allowing the complete installation and setup to be handled automatically by Snakemake.

LysoPhD does not appear to be available to download anywhere and was dropped from the comparison. ProphET requires the unsupported BLAST legacy and EMBOSS packages. It is not available on any package manager and instructions for a clean installation are incomplete and not compatible with conda. The codebase was last updated in 2019. Numerous issues were encountered installing dependencies and despite significant effort we were not able to create a working installation. ProphET’s installation script reported many errors during setup, but alarmingly finished with an exit code zero to indicate a *successful* installation. Preparing the necessary GFF files in a format that the program could use was non-trivial. The program reported errors during runtime that we believe are related to the errors encountered during installation; ProphET terminated with incomplete output but again returned an exit code zero to indicate a *successful* run. ProphET was dropped from the comparison.

### Prophage prediction performance

There was minimal difference in the performance metrics for the different methods of running PhiSpy, and we have recently shown (Roach et al in preparation) that including HMM searches with PhiSpy results in less than one additional prophage being identified. Therefore, only PhiSpy using default settings will be discussed in comparison to the other tools. PhiSpy, VIBRANT, and Phigaro performed best for mean accuracy (Figure 1a; Table S3) while DBSCAN-SWA performed the worst. PhiSpy, Phigaro, and Phage Finder performed best for mean precision (Figure 1b; Table S3). DBSCAN-SWA, PhageBoost, VirSorter, and VirSorter2 all performed poorly for mean precision. This was mostly driven by a high false-positive rate compared to the other tools (Figure S1). PhiSpy, VirSorter, VirSorter2, VIBRANT, DBSCAN-SWA and PhageBoost all had high mean recall scores.

**Figure 1:**
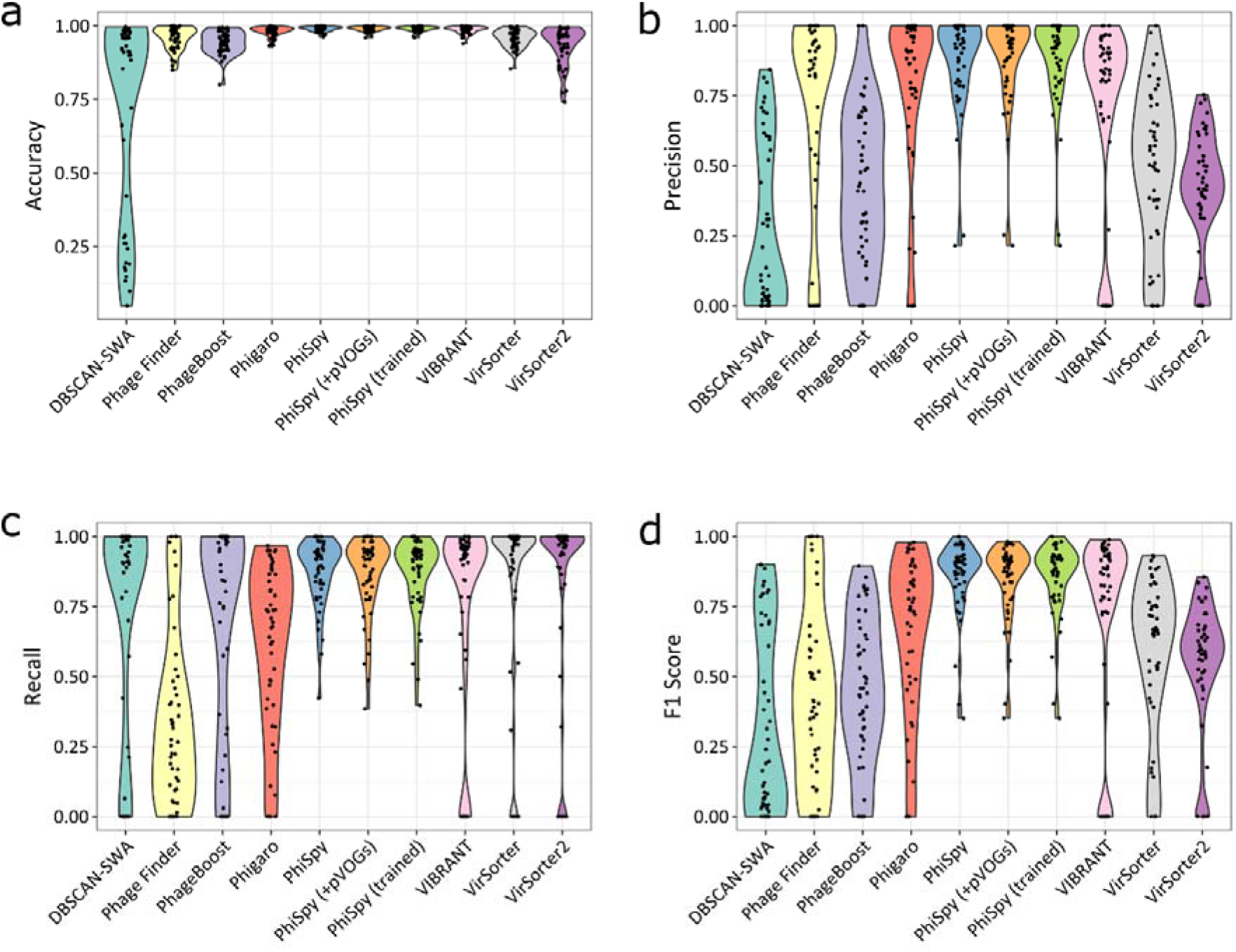
Prediction performance metrics for prophage callers. Violin plots for each tool are shown with individual points for each genome indicated. The graphs show: ‘Accuracy’ (*a*) as the ratio of correctly labelled genes to all genes, ‘Precision’ (*b*) as the ratio of correctly labelled phage genes to all predicted phage genes, ‘Recall’ (*c*) as the ratio of correctly labelled phage genes to all known phage genes, and ‘f1 Score’ (*d*) as defined in the methods. For all graphs, more is generally better.

Each tool balances between recall and precision. For example, the more conservative Phage Finder performed relatively well in terms of precision, making very confident predictions, but had one of the lower mean recall ratios and was not predicting prophages based on limited information. In contrast, the more speculative DBSCAN-SWA and PhageBoost both exhibited the opposite trend.

The f_1_ Score is a more nuanced metric, as it requires high performance in both precision and recall. PhiSpy, VIBRANT, Phigaro, VirSorter, and VirSorter2 all averaged above 0.5, while the remaining tools suffered from too many false predictions (FP or FN) (Figure 1d).

### Runtime performance

Many users will not be too concerned about runtime performance, for instance if they are performing a one-off analysis on a genome of interest all the tools will finish in a reasonable time. However, efficient resource utilization is an important consideration for large-scale analyses. Provisioning computing resources costs money and a well optimised tool that runs fast translates to real-world savings. The runtime distributions across the genomes are shown for each tool in Figure 2a. The slowest prophage predictors were generally VirSorter and VirSorter2 with mean runtimes of 1,316 and 2,118 seconds respectively, except for a single DBSCAN-SWA run taking 4,697 seconds. PhiSpy using the trained datasets was by far the fastest performing tool (8.4 seconds mean runtime), although if an appropriate training set is not available for the genus of interest it would first need to be generated to benefit from these reduced runtimes. PhageBoost was the next fastest (37.8 seconds mean runtime) and Phage Finder, Phigaro, and PhiSpy with default parameters all performed similarly well in terms of runtime.

**Figure 2:**
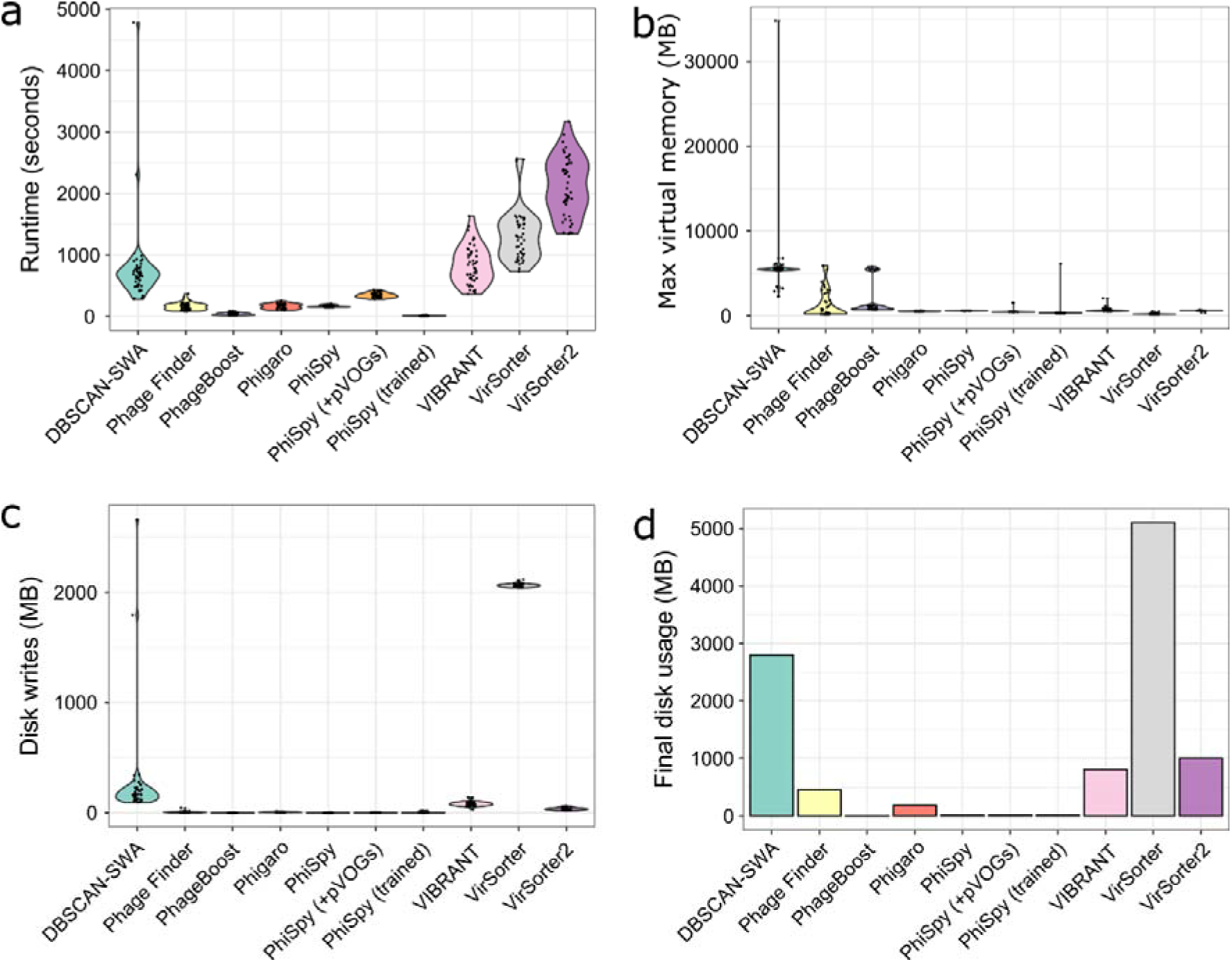
Runtime and peak memory usage comparison. Violin plots for each tool are shown with individual points for each genome indicated. The graphs show total runtime in seconds (*a*), peak memory usage in MB (*b*), total file writes in MB (*c*) and the final total disk usage (all genomes) in MB (*d*). For all graphs, less is better.

Memory requirements also remain an important consideration for provisioning resources for large-scale analyses. For instance, inefficiency is encountered where the memory required by single-threaded processes exceeds the available memory per CPU. Peak memory usage for each tool is shown in Figure 2b. Memory requirements were lowest for VirSorter and trained PhiSpy with 210 and 450 MB mean peak memory respectively. There was a single notable exception for trained PhiSpy (predicting prophages in *E. coli* O157:57 EDL933) with a peak memory usage of 6.13 GB. DBSCAN-SWA had the highest mean peak memory of 6.0 GB with one run requiring 35 GB at its peak. Apart from the DBSCAN-SWA outlier, there were no situations where the peak memory usage would prevent the analysis from completing on a modest personal computer, but at larger-scales, Phigaro, PhiSpy, VirSorter, and VirSorter2 have an advantage in terms of peak memory usage.

Another important consideration for large-scale analyses are the file sizes that are generated by the different tools. Large output file sizes can place considerable strain on storage capacities, and large numbers of read and write operations can severely impact the performance of a system or HPC cluster for all users. Total file writes for the default files (in MB, including temporary files) are shown in Figure 2c and the final disk usage for all genomes for each tool is shown in Figure 2d. VirSorter, DBSCAN-SWA, and VirSorter2 performed the most write operations with mean file writes of 2.063, 0.262, and 0.034 GB respectively. The other tools performed similarly well and have a clear advantage at scale as they perform far fewer disk writes. VirSorter and DBSCAN-SWA removed most of their generated files, however, the final disk usage for these tools were still the highest at 5.36 and 2.96 GB respectively. Disk usage for PhageBoost and PhiSpy was by far the lowest at 0.14 and 15 MB respectively.

## Caveats

Every bioinformatics comparison involves many biases. In this comparison, PhiSpy performs well, but we developed PhiSpy and many of the gold-standard genomes were extensively used during its development to optimize the algorithm. VirSorter and VirSorter2 were primarily developed to identify viral regions in metagenomes rather than prophages in bacterial genomes—although they have been used for that e.g. in Glickman et al. (2020)—and filtering VirSorter and VirSorter2 hits with CheckV (Nayfach et al., 2021) is recommended. By openly providing the Prophage Prediction Comparison framework, creating a framework to install and test different software, and defining a straightforward approach to labelling prophages in GenBank files, we hope to expand our gold-standard set of genomes and mitigate many of our biases. We welcome the addition of other genomes (especially from beyond the Proteobacteria/Bacteroidetes/Firmicutes that are overrepresented in our gold-standard database).

Recent developments in alternative approaches to predict prophages, including mining phage-like genes from metagenomes and then mapping them to complete genomes (Nayfach et al., 2021) and using short-read mapping to predict prophage regions from complete bacterial genomes (Kieft and Anantharaman, 2021) have the potential to generate many more ground-truth prophage observations. However, both approaches are limited as they will identify prophages that are active, but are unable to identify quiescent prophage regions, and thus for prophage prediction algorithms they will provide useful true positive datasets but may not provide accurate true negative datasets.

## Conclusions

In this comparison, PhiSpy, VIBRANT, and Phigaro were the best performing prophage prediction tools for f_1_ score. PhiSpy and Phigaro were also among the best in terms of runtime performance metrics. Phage Finder performs well in terms of precision at the expense of false-negatives, whereas VirSorter, VirSorter2, DBSCAN-SWA and PhageBoost perform well for recall at the expense of false-positives. Currently, DBSCAN-SWA, VirSorter, and VirSorter2 are not as well suited for large-scale identification of prophages from complete bacterial genomes when compared to the other tools. More genomes with manually curated prophage annotations are needed, and we anticipate that these benchmarks will change with the addition of new genomes, the addition of new tools, and as the tools are updated over time. Developers are strongly encouraged to contribute by adding or updating their tool and adding their manually curated genomes to be included in the benchmarking. Users are strongly encouraged to check the GitHub repository for the latest results before making any decisions on which prophage prediction tool would best suit their needs.

## Author contributions

RAE conceived of the study; KM and PD generated the initial gold-standard set and SKG, LI, and EP contributed to the gold-standard set; RAE and MJR created the framework; RAE, MJR, and SR performed the analysis. All authors contributed to the manuscript writing.

## Acknowledgments

This work supported by the National Institute Of Diabetes And Digestive And Kidney Diseases of the National Institutes of Health under Award Number RC2DK116713 to RAE. The support provided by Flinders University for HPC research resources is acknowledged.

## Data availability

All the data is available at DOI: 10.5281/zenodo.4739878 and from https://github.com/linsalrob/ProphagePredictionComparisons/tree/v0.1-beta

## Supplementary data

**Table S1. Genomes provided in the gold-standard library with manually curated prophages**

**Table S2. Prophages identified in the genomes**

**Table S3.** Mean metrics for each tool as measured from our gold-standard set of genomes.

| *Tool* | *Accuracy* | *Precision* | *Recall* | *f<sub>1</sub> score* |
| --- | --- | --- | --- | --- |
| DBSCAN-SWA | 0.72 | 0.30 | 0.72 | 0.33 |
| Phage Finder | 0.95 | 0.76 | 0.35 | 0.43 |
| PhageBoost | 0.94 | 0.45 | 0.70 | 0.45 |
| Phigaro | 0.98 | 0.82 | 0.61 | 0.65 |
| PhiSpy | 0.99 | 0.88 | 0.87 | 0.85 |
| VIBRANT | 0.99 | 0.70 | 0.75 | 0.72 |
| VirSorter | 0.96 | 0.49 | 0.83 | 0.58 |
| VirSorter2 | 0.93 | 0.42 | 0.82 | 0.54 |

**Figure S1.**
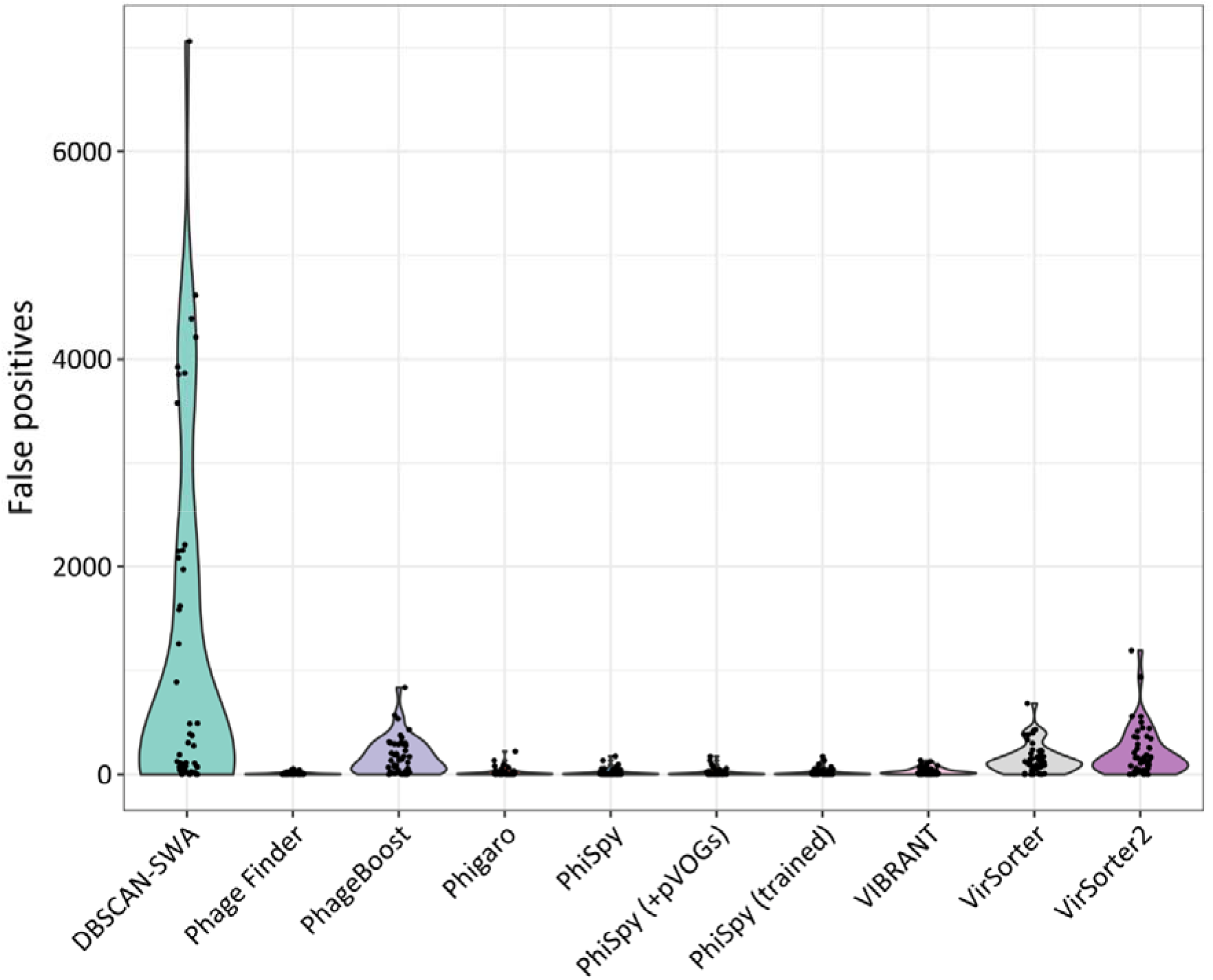
False positive comparison. Violin plots for each tool show ‘False Positives’ as the number of genes incorrectly labelled prophage genes in each genome. Less is better.

