## Supplementary figures and images for "Philympics 2021: Prophage Predictions Perplex Programs"

### Figure S1

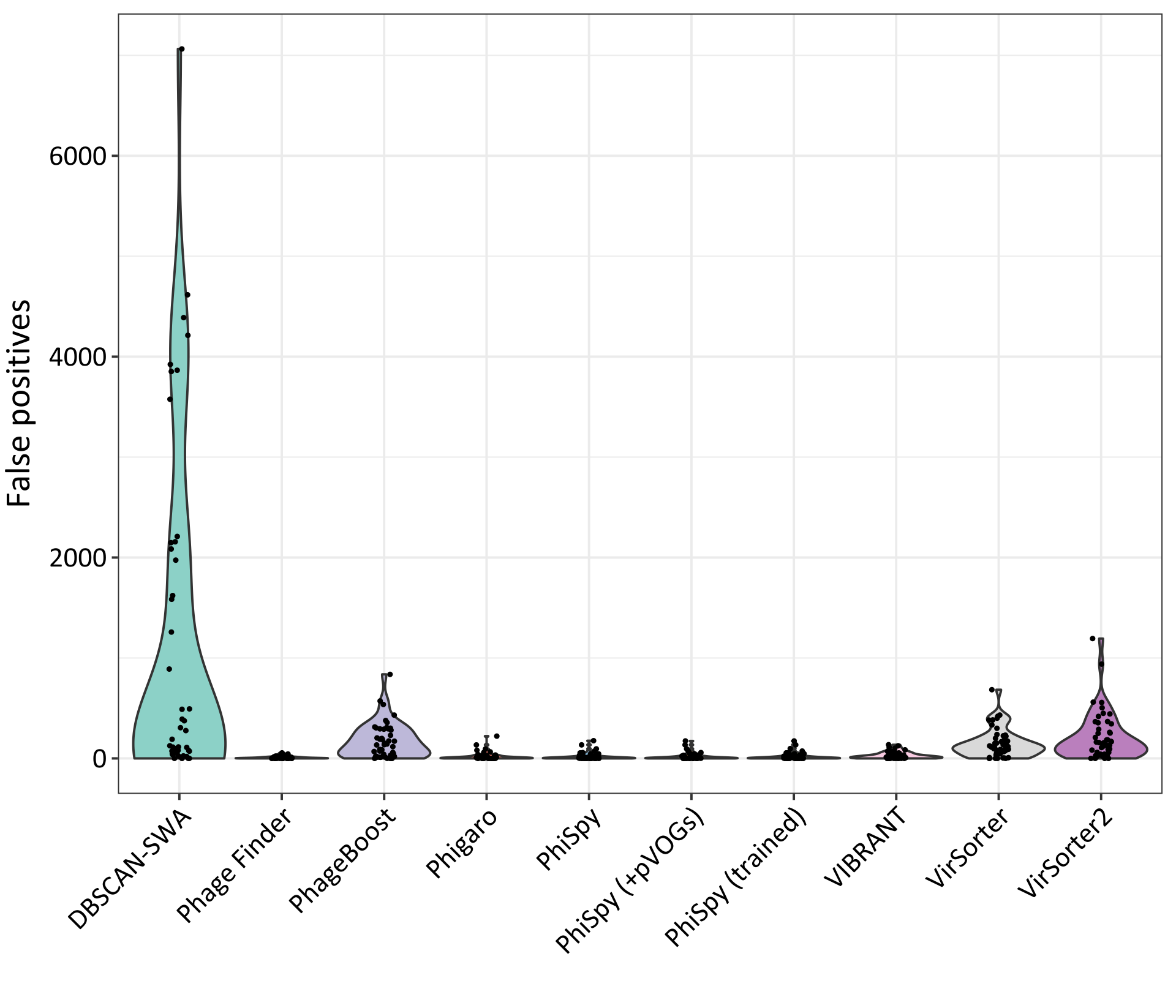
